# Blood flow-induced Notch activation and endothelial migration enable embryonic vascular remodeling

**DOI:** 10.1101/095307

**Authors:** Bart Weijts, Edgar Gutierrez, Semion K Saikin, Ararat J Ablooglu, David Traver, Alex Groisman, Eugene Tkachenko

## Abstract

Arteries and veins are formed independently by different types of endothelial cells (ECs). In vascular remodeling, arteries and veins become connected and some arteries become veins. It is unclear how ECs in transforming vessels change their type and how fates of individual vessels are determined. In embryonic trunk, vascular remodeling transforms arterial intersegmental vessels (ISVs) into a functional network of arteries and veins. We found that, once an ISV is connected to venous circulation, venous blood flow promotes upstream migration of ECs that results in displacement of arterial ECs by venous ECs, completing the transformation of this ISV into a vein without trans-differentiation of ECs. Arterial blood flow initiated in two neighboring ISVs prevents their transformation into veins by activating Notch signaling in ECs. Together, different responses of ECs to arterial and venous blood flow lead to the formation of a balanced network with equal numbers of arteries and veins.

## INTRODUCTION

Blood vessels remodel in response to varying conditions and functional demands during development, tissue regeneration and organ growth. Failure in vascular remodeling can lead to ischemia, an inadequate blood supply to an organ. There is compelling evidence that vascular remodeling does not occur without blood flow^1,2^. However, the mechanisms and processes underlying the sensing of blood flow and responses to it in the context of vascular remodeling are poorly understood.

One of the paradigms of vascular remodeling is the formation of a functional network of intersegmental vessels (ISVs) in the trunk of zebrafish embryo^3,4^. Initially, all ISVs are arterial with no blood flow, and the remodeling transforms them into a balanced network with nearly equal numbers of arteries and veins. The transformation of an arterial ISV into venous starts when a venous sprout anastomoses with the ISV, connecting it to the posterior cardinal vein (PCV). While anatomically a vein, this ISV is still lined with arterial endothelial cells (ECs), which differ from venous ECs in multiple respects^5–7^. The establishment of venous identity of this ISV is completed after ECs lining it change to venous (Fig. 1a). This change is believed to occur by trans-differentiation of arterial ECs into venous ECs. It remains unclear, however, how fates of individual ISVs are determined, and what mechanisms lead to a strong preference (70-80%) of venous ISVs to be flanked by arterial ISVs^8^.

**Figure 1:**
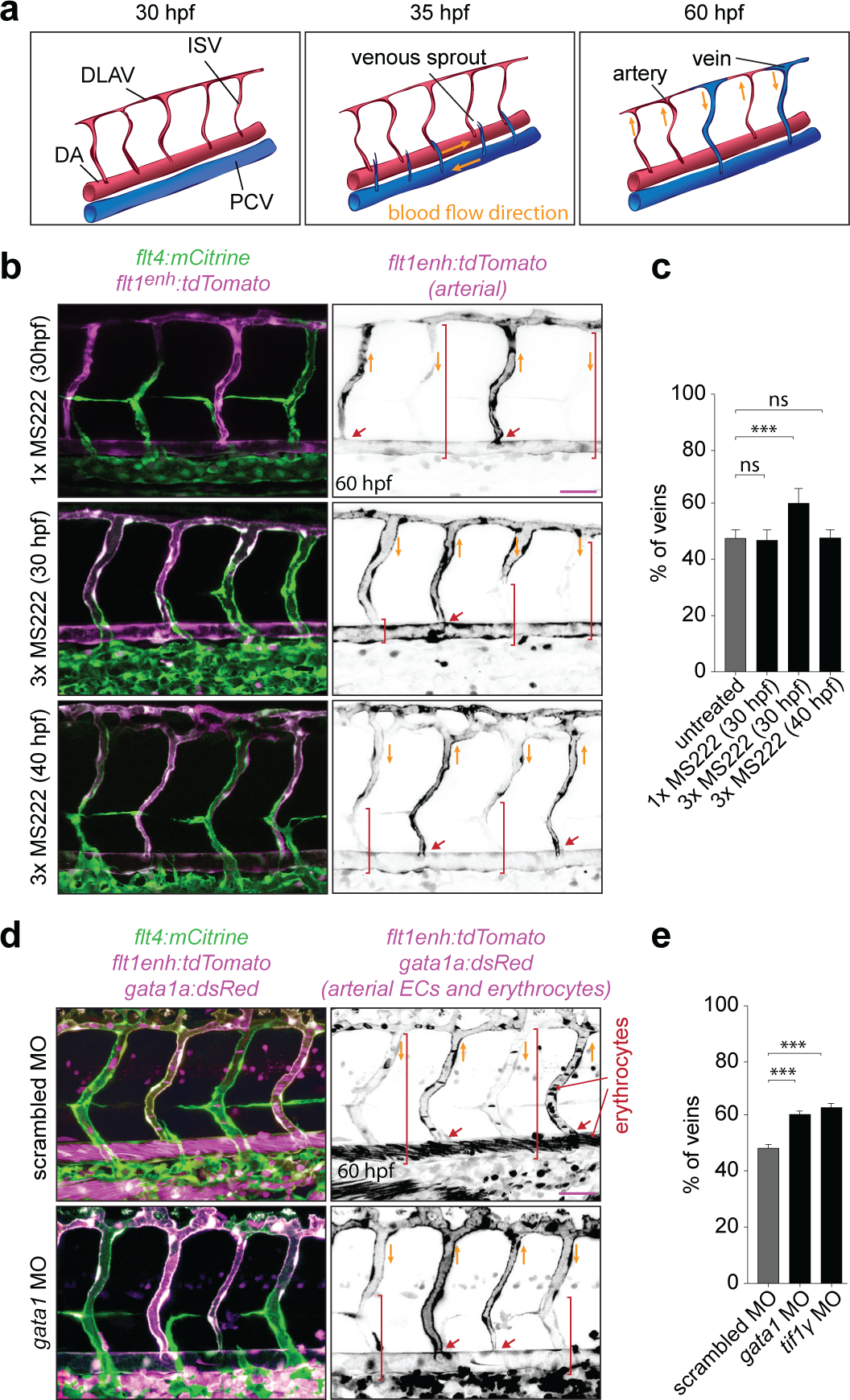
Blood flow controls vascular remodeling of the trunk. Panels in (**b**) and (**d**) show lateral images of zebrafish embryos at 60 hpf with anterior side facing left. Venous ECs are labelled with mCitrine and arterial ECs are labelled with mCitrine and tdTomato. *Orange arrows* indicate the direction of blood flow, *red arrows* point to arterial ISVs and *red brackets* highlight regions of venous ISVs without arterial ECs. Scale bars are 25 μm. The numbers are averages ± SEM from at least three independent experiments with a minimum of n=25 animals per conditions per experiment. ‘***’ indicates P<0.001. **a**) Schematic overview of intersegmental vasculature remodeling. DA - dorsal aorta, PCV - posterior cardinal vein, DLAV - dorsal longitudinal anastomotic vessel, ISV-intersegmental vessel. **b**) Flow of blood was reduced by administering 3x MS222 at indicated time point. **c**) Percentage of venous ISVs in embryos with reduces blood flow. **d**) Viscosity of blood was reduced by morpholino knock down of *gatala* or *tiflγ* which are required for the formation of erythrocytes. Erythrocytes are marked by dsRed. **e**) Percentage of venous ISVs in embryos without erythrocytes.

## RESULTS

### Blood flow controls vascular remodeling of the trunk

To investigate the role of blood flow in the remodeling of the intersegmental vasculature, we employed transgenic zebrafish in which arterial and venous ECs are differently labeled with fluorescent markers^8^. We treated 30 hpf embryos with muscle relaxant (ms-222; tricaine methanesulfonate), which lowers the heart rate, thus reducing the blood flow. The proportion of venous ISVs at 60 hpf increased to significantly greater than 50% (Fig. 1b,c; Extended Data Fig. 1a,b). Moreover, these venous ISVs were lined with venous ECs only in ventral parts, whereas their dorsal parts remained lined with arterial ECs (Fig. 1b). To test whether these remodeling defects were related to a reduction of flow shear stress at vessel walls (which is known to be sensed by ECs) caused by reduced heart rate, we suppressed the formation of erythrocytes through morpholino oligonucleotide (MO) mediated knock down of *gatal*^9^ or *tifly*^10^. The heart rate in these embryos was 1.7 times higher and the blood flow rate was 1.9 times higher than normal (Extended Data Fig. 1c).

Nevertheless, based on ~3 times greater viscosity of blood as compared with plasma, as measured in large vessels11, the vessel wall shear stress in embryos without erythrocytes was expected to be ~1.5 times lower. Moreover, the effective viscosity of blood in small vessels (with the diameter comparable to the size of erythrocytes) has been reported as >6 times greater than the viscosity of plasma^12,13^, suggesting a >3-fold reduction in shear stress in ISVs of embryos without erythrocytes. These embryos had vascular remodeling defects similar to those in the embryos with reduced heart rate (Fig.1d,e). To distinguish the effect of blood flow on the proportion of venous ISVs and on the type of ECs lining venous ISVs, we reduced blood flow after the fate determination of ISVs (arterial vs. venous) was largely complete (at 40 hpf). Whereas the proportion of venous ISVs in these embryos was normal (50%), venous ISVs were still defective in terms of their EC lining (Fig. 1b,c). Taken together, our results indicate that blood flow controls the proportion of venous ISVs and, in addition, is required for the ECs lining the venous ISVs to change from arterial to venous.

### Blood flow-dependent displacement of arterial ECs by venous ECs in venous ISVs

To find out how the type of ECs in ISVs changes from arterial to venous, we used zebrafish embryos in which arterial ECs and erythrocytes express red fluorescent proteins and all ECs express yellow fluorescent protein. Once blood flow through a newly formed venous ISV was initiated, ECs started migrating against the flow, and this migration eventually resulted in the displacement of arterial ECs by venous ECs from the PCV (Fig. 2a, Extended Data Fig. 2a and Supplementary Video 1,2). During the displacement, some venous ECs divided (Extended Data Fig. 2b and Supplementary Video 3). Lineage tracing expFriments confirmed that ECs lining venous ISVs at the end of remodeling originated from the PCV, while ECs originally lining these ISVs had migrated into the dorsal longitudinal anastomotic vessel (Fig. 2c). In contrast, we found no evidence of EC migration in arterial ISVs (Fig. 2b,d and Supplementary Video 4). Thus, our results indicate that the change in the type of ECs in venous ISVs occurs through blood flow-dependent EC migration and displacement of arterial ECs by venous ECs.

**Figure 2:**
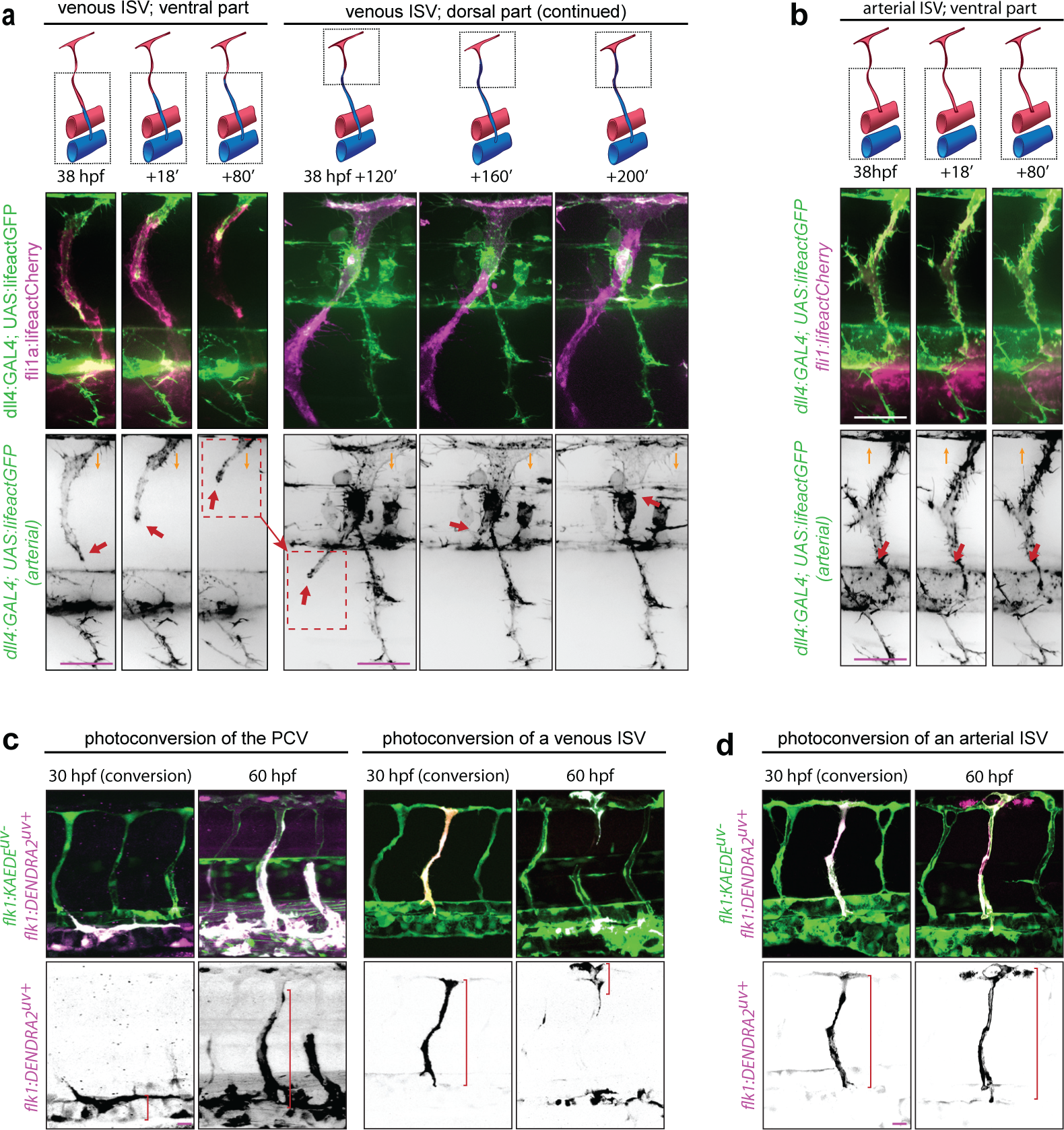
Blood flow-dependent displacement of arterial ECs by venous ECs in venous ISVs. Lateral images of zebrafish embryos with anterior side facing left. *Orange arrows* indicate the direction of blood flow through the ISVs. Scale bars are 25 μm. **a,b**) Venous ECs are labelled with lifeactCherry and arterial ECs are labelled with lifeactGFP and lifeactCherry. **a**) Stills from Video 2. *Red arrows* point at an arterial EC migrating in a venous ISV. **b**)Stills from Video 4. *Red arrows* point at an arterial EC in an arterial ISV. **c,d**) All ECs express the photoconvertible (green-to-red) fluorescent protein DENDRA2. *Red brackets* highlight ECs with photo-converted DENDRA2. The photo-conversion was done at 30 hpf in the posterior cardinal vein (PCV) (*left panel **c***), a venous ISV (*right panel* **c**) or an arterial ISV (**d**).

### Blood flow promotes EC migration in veins but not in arteries

Arterial ECs have been shown to migrate against the flow in some arteries^14,15^, but we found arterial ECs migrating in venous but not arterial ISVs. Planar polarity of cells often correlates with the direction of cell migration^16^, and, in agreement with a previous report^17^, we found ECs polarized against blood flow in both venous and arterial ISVs (Extended Data Fig. 3a, b). Moreover, we have previously shown that the level of EC planar polarization correlates with the flow shear stress^18^. To test whether shear flow can induce EC migration, we performed *in vitro* experiments in microfluidic perfusion chambers, where we applied a range of shear stresses to confluent human umbilical venous endothelial cells (HUVECs) and human umbilical arterial endothelial cells (HUAECs). After the flow was initiated, HUVEC gradually became polarized against the flow and started to migrate against the flow, with the onsets of polarization and directed migration occurring nearly simultaneously (Extended Data Fig. 3c; Supplementary Video 5). Furthermore, the time it took HUVECs to start migrating against the flow decreased with shear stress, whereas the ultimate migration velocity increased with shear stress (Fig. 3a, b; Supplementary Video 6). HUAECs also migrated against the flow at sufficiently high shear stress (Extended Data Fig. 3d). Together, our *in vitro* experiments show that both arterial and venous ECs polarize and migrate against sufficiently strong shear flow.

**Figure 3:**
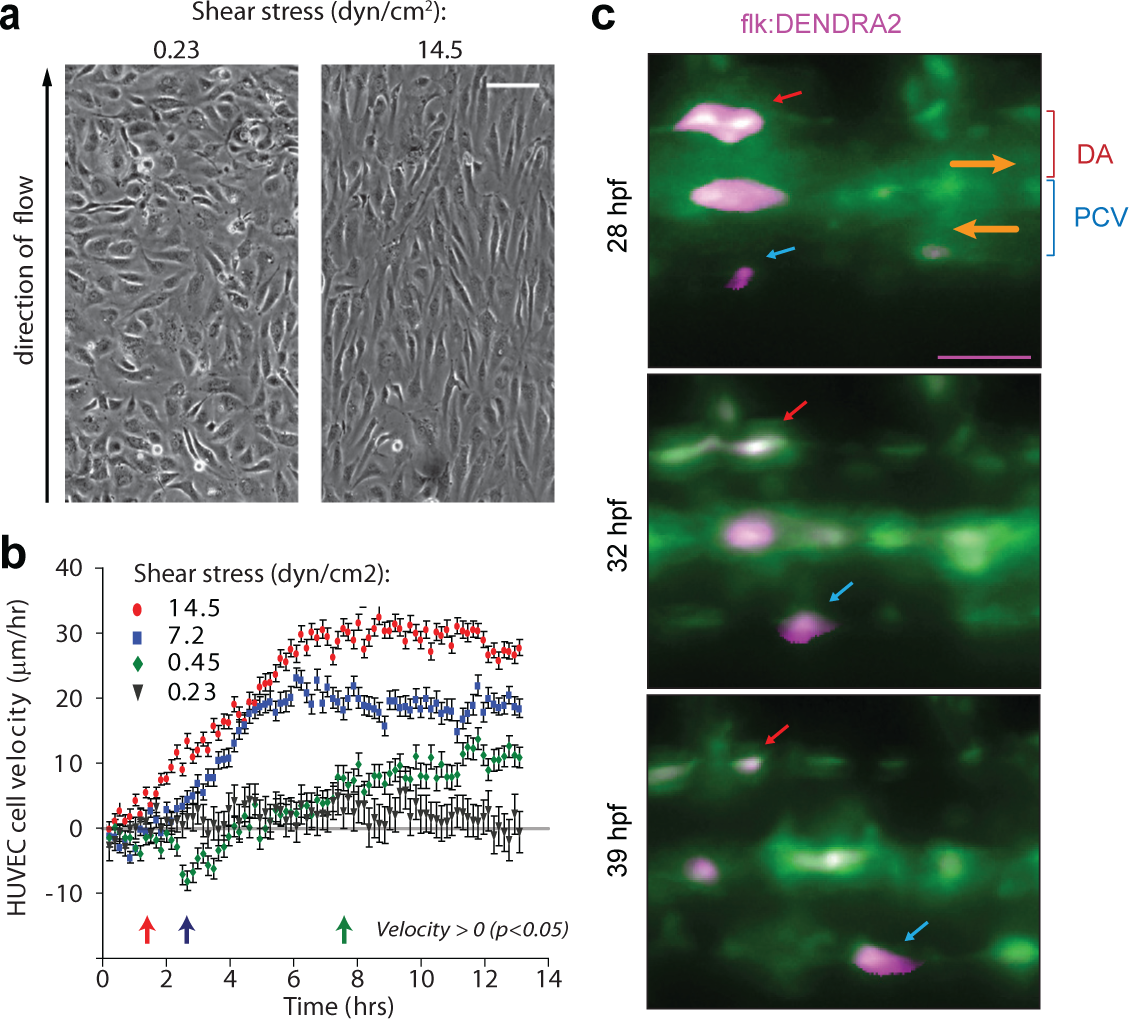
Blood flow promotes EC migration in veins but not in arteries. **a**) Phase images from Supplementary Video 6. showing confluent HUVECs after 10 hrs under laminar flow with shear stresses of 0.23 or 14.5 dyn/ cm2. Scale bar is 100 μm. **b**) Average velocities of upstream migration for HUVECs exposed to different shear stresses as functions of time after the inception of shear flow (n=250 to 600 for individual shear stresses). Arrows at the bottom (colors correspond to those of the velocity data points) indicate the time points at which the migration of cells against the flow becomes statistically significant (average upstream velocity becomes positive with p<0.05). **c**) All ECs express photo-convertible (green-to-red) fluorescent protein DENDRA2. The photo-conversion was done at 28 hpf in the DA and PCV. Lateral images of zebrafish embryo with anterior facing left. *Orange arrows* indicate the direction of blood flow. *Red arrows* point to an arterial EC and *blue arrows* point to a venous EC. Scale bar is 25 μm.

Arterial and venous blood flow have different dynamics, and these differences may affect EC migration. We found that blood flow in arterial ISVs was strongly pulsatile, with minimum diastolic velocity close to 0, peak systolic velocity ~ 1500 μm/sec, the amplitude of velocity variations nearly equal to the mean velocity, and characteristic rate of velocity variation at ~ 7500 μm/sec2. The pulsations in venous ISVs were much weaker, with velocity varying between ~250 to ~750 μm/ sec (Extended Data Fig. 3e; Supplementary Video 7), corresponding to 1:2 ratio between the amplitude of variations and mean velocity, and characteristic rate of velocity change at ~ 2500 μm/ sec^2^. The dynamics of blood flow in the dorsal aorta (DA) resemble those in arterial ISVs, whereas the dynamics of blood flow in the PCV are similar to those in venous ISVs (Extended Data Fig. 3e,f; Supplemental Video 8). We observed migration of ECs against blood flow in the PCV and no net migration in the DA, very much in line with EC migration in venous and arterial ISVs (Fig. 3c; Extended Data Fig. 3g-j). Together, these results can be explained by an assumption that strong pulsation of blood flow suppresses the upstream migration of ECs.

### Arterial blood flow protects ISVs from transforming into veins by activating Notch signaling

Previous studies showed that inhibition of Notch signaling through abrogation of its main endothelial receptor *delta like 4* (*dll4*) leads to a greater proportion of arterial ISVs being transformed into venous ISVs^19,20^. It has also been shown that Notch activity is regulated by blood flow^21–23^. When we inhibited Notch signaling, we observed a phenotype similar to that caused by reduced shear stress, in terms of both an increased proportion of venous ISVs and incomplete replacement of arterial ECs with venous ECs in venous ISVs (Fig. 1b, 4f; Extended Data Fig. 4a,b). We visualized the effect of blood flow on Notch signaling in ECs in ISVs by imaging transgenic embryos *Tg*(*Tp1:d2GFP*)^24^ that carry a Notch signaling reporter expressing a destabilized GFP (d2GFP) and found that the initiation of blood flow through ISVs enhanced the expression of d2GFP in ECs in arterial ISVs but not in other vessels (Fig. 4a,b; Supplementary Video 9,10). Furthermore, we observed that venous sprouts did not anastomose with arterial ISVs with enhanced expression of d2GFP (Fig. 4a,b). Embryos with reduced blood flow had significantly lower expression of d2GFP in arterial ISVs, indicating that the level of Notch signaling in arterial ISVs depends on the strength of blood flow (Fig. 4c; Extended Data Fig. 4c). The level of d2GFP expression prior to the initiation of blood flow varied between ISVs, but these variations in the early expression of d2GFP did not seem to correlate with the subsequent fate determination of ISVs (Extended Data Fig. 4d). The early expression of d2GFP usually subsided to undetectable levels soon after ISVs became anatomical veins (Extended Data Fig. 4d; Supplementary Video 11). Together, these results suggest that Notch signaling is activated by arterial but not venous blood flow, and that this activation protects arterial ISVs from anastomosis with venous sprouts.

**Figure 4:**
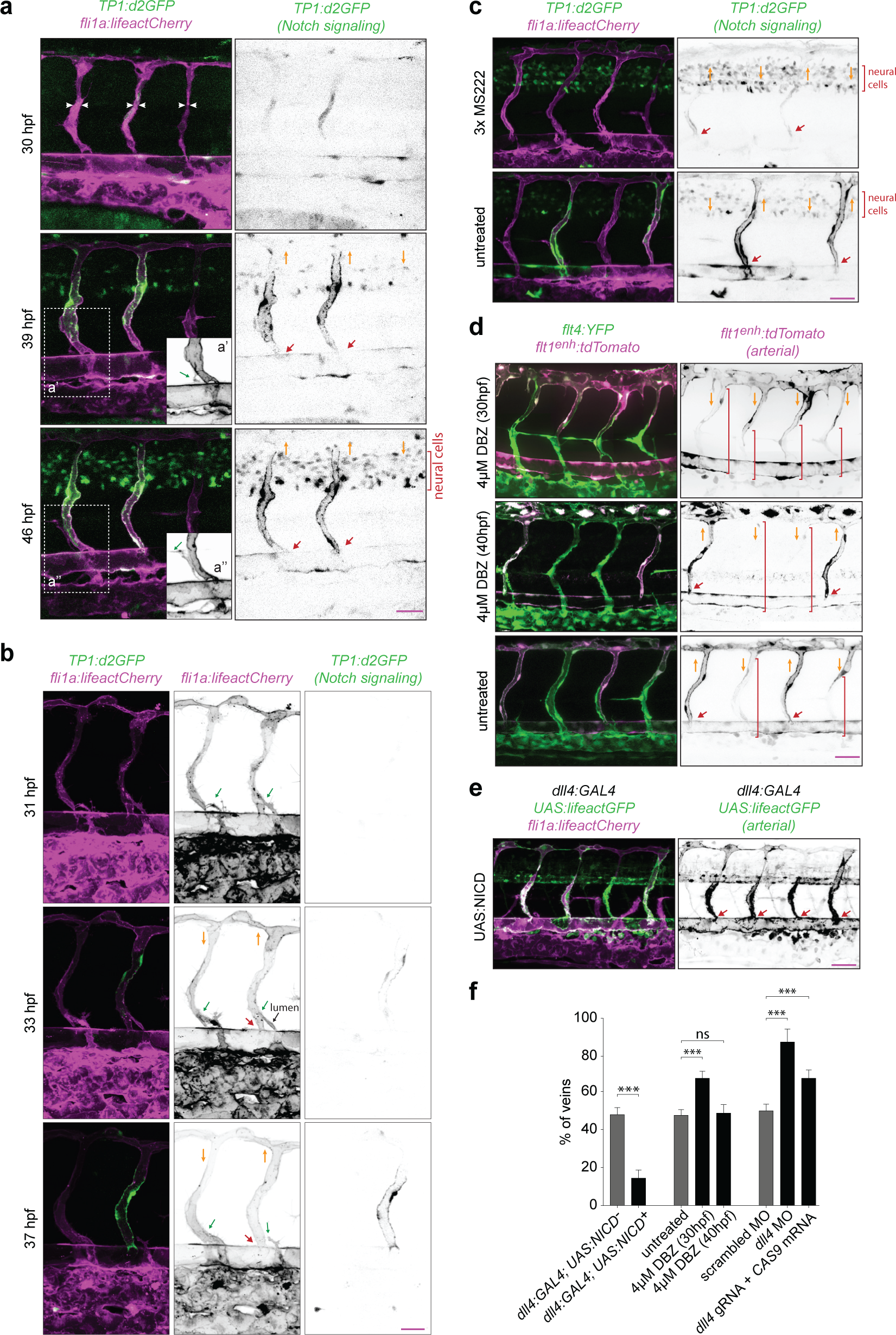
Arterial blood flow protects ISVs from transforming into veins by activating Notch signaling. Panels (**a**),(**b**),(**c**),(**d**) and (**e**) show lateral images of zebrafish embryos with anterior side facing left. *Red arrows* point to arterial ISVs. *Orange arrows* indicate the direction of blood flow. For panels (**b**),(**d**) and (**e**) images were acquired at 60 hpf. Scale bars are 25 μm. The numbers are averages ± SEM. ‘***’ indicates P<0.001. **a, b**) Notch signaling is reported by the expression of destabilized GFP (d2GFP) under the control of 12xCSL Notch responsive elements. **a**) Stills from Video 9. *White arrowheads* highlight ISVs without blood flow. Insets **a**’ and **a**” show expression of lifeactCherry (grey). *Green arrows* in the insets point to a venous sprout that did not anastomose with an arterial ISV. **b**) Stills from Video 11. *Green arrows* point to venous sprouts. The sprout on the right detached from an ISV expressing d2GFP before a functional connection with the PCV was formed, as indicated by the extent of lumen formation (*black arrow*). **c**) Expression of Notch signaling reporter in embryos with reduced blood flow. **d**) Inhibition of Notch signaling at different time points of the development. Venous ECs are labelled with mCitrine and arterial ECs are labelled with mCitrine and tdTomato. *Red brackets* highlight regions of venous ISVs without arterial ECs. **d**) Ectopic expression of the Notch Intracellular Domain (NICD) specifically in arterial ECs. Venous ECs are labelled with lifeactCherry and arterial ECs are labelled with lifeactGFP and lifeactCherry. **f**) The percentage of venous ISVs in embryos with perturbed Notch signaling. (At least three independent experiments with a minimum of n=25 animals per conditions per experiment).

To further test the role of Notch signaling in the protection of arterial ISVs from transformation into veins^19,20^, we inhibited Notch signaling at 30 hpf, shortly before anastomosis of arterial ISVs with venous sprouts was about to start. To this end, we administered a γ-secretase inhibitor dibenzazepine (DBZ), which prevented the cleavage and release of transcriptionally active Notch intracellular domain (NICD). DBZ treatment resulted in a significantly larger proportion of venous ISVs (Fig. 4d,f), indicating that Notch signaling is required for the protection of arterial ISVs from transformation into veins. Last but not least, constitutive activation of Notch signaling in arterial ECs protected nearly all ISVs from transformation into veins (Fig. 4e,f). Therefore, our results confirmed that Notch signaling protects arterial ISVs from transformation into veins.

In embryos in which Notch inhibition resulted a greater proportion of venous ISVs, the flow of blood through venous ISVs was slower (as judged by the tracking of erythrocytes), and venous ISVs had arterial ECs in their dorsal parts, the phenotype resembling the effect of reduced blood flow (Fig. 1b, 4d). To test whether Notch signaling has a direct effect on the change in the type of ECs in venous ISVs, we inhibited Notch signaling after the fate determination of ISVs was largely complete (at 40 hpf). Despite the Notch inhibition, the displacement of arterial ECs by venous ECs in venous ISVs proceeded normally (Fig. 4d). Moreover, there was no directed migration of ECs in arterial ISVs (Extended Data Fig. 4e), and there were no other obvious defects in the vascular remodeling (Fig. 4f; Extended Data Fig. 4f-i). Together, our results indicate that blood flow-induced Notch signaling protects arterial ISVs from transformation into venous ISVs, but does not have a major direct effect on EC migration in response to blood flow.

## Discussion

Our results can be explained by the following model (Fig. 5). After a venous sprout originating from the PCV anastomoses with an ISV, venous blood flow through this ISV and arterial blood flow through the two immediately adjacent ISVs are initiated. Arterial blood flow activates Notch signaling in ECs in these adjacent ISVs, thereby preventing their anastomosis with venous sprouts and their transformation into veins. On the other hand, the arterial blood flow through ISVs that are next to the two immediately adjacent ones is too weak to prevent their anastomosis with venous sprouts. As a result, by the end of the remodeling, venous ISVs are flanked by arterial ISVs, arterial ISVs are flanked by venous ISVs, and the numbers of venous and arterial ISVs are nearly equal, leading to efficient blood circulation. In ISVs connected to the PCV, venous blood flow induces upstream migration of ECs, leading to the displacement of arterial ECs by venous ECs. In contrast, in arterial ISVs, arterial ECs do not migrate (likely, because of the highly pulsatile character of blood flow), and venous ECs do not displace them. Thus, responses of arterial ECs to arterial blood flow provide a feedback mechanism ensuring robust formation of a functional ISV network, while responses of both arterial and venous ECs to venous blood flow lead to the change of the type of ECs in venous ISVs.

**Figure 5:**
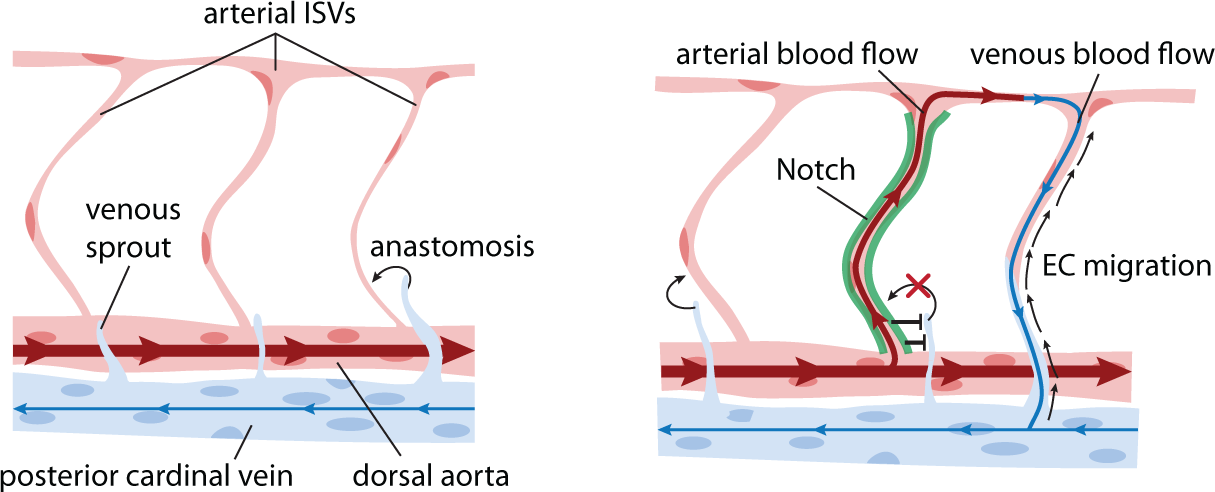
Model explaining how blood flow-induced Notch signaling and endothelial migration enable the remodeling of all-arterial intersegmental vasculature into a network with ~1:1 ratio of arteries and veins.

With an exception of the first four and the last two ISVs, the ISVs have no bias toward either arterial or venous fate^48^. In the proposed model, the fate of an ISV depends on the time when a venous sprout reaches this ISV^4^. An early sprout is likely to reach an ISV without blood flow and anastomose with it, initiating its transformation into a vein, whereas a late sprout is likely to reach an ISV that is protected from anastomosis by arterial blood flow. Sprouts that are unable to anastomose with ISVs continue to grow dorsally to become part of the lymphatic vasculature. Recently, it has been suggested that venous sprouts expressing *prox1* at their emergence are pre-destined to become part of the lymphatic system^25^. However, we found that the inhibition of Notch signaling resulted in the transformation of nearly all ISVs into anatomical veins, suggesting that each venous sprout has a potential to anastomose with an ISV. The bias of *prox1* positive venous sprouts towards the lymphatic fate may be due to their relatively late outgrowth.

We found that in ISVs with arterial ECs and arterial blood flow, ECs do not migrate and have high Notch activity. In contrast, in ISVs with venous blood flow, there is no Notch activation in arterial ECs and both arterial and venous ECs migrate upstream. Surprisingly, the effect of the type of blood flow on EC migration does not appear to depend on Notch signaling. It is unclear what parameters of arterial blood flow allow ECs to distinguish it from venous flow. Plausible candidates are the maximal shear stress, the ratio between the maximal and mean (or minimal) stress, and the rate of change of shear stress (which is proportional to the rate of change in blood flow velocity).

Embryonic ECs differentiate into arterial and venous before blood vessels start forming^26–29^. The maintenance of EC identity is important for arterial and venous angiogenesis^30–32^. For example, in zebrafish trunk, arterial ECs sprout in response to Vegfa, whereas venous ECs sprout in response to Vegfc and Bmp^33–35^. On the other hand, ECs are believed to have plasticity, enabling transdifferentiation between arterial and venous ECs in response to certain environmental cues^36,37^. Given this context, the finding of this study that arterial ECs in venous ISVs do not trans-differentiate into venous ECs but are replaced by venous ECs migrating from the PCV is intriguing. It shows that plasticity of ECs is not necessary for plasticity of blood vessels.

### Materials and Methods *Zebrafish husbandry*

Zebrafish (*Danio rerio*) were maintained according to the guidelines of the UCSD Institutional Animal Care and Use Committee. The following zebrafish lines have been previously described: *Tg(fli1a:GFP*^*y1 38*^, *Tg*(*UAS:lifeactGFP*)^*mu27139*^, *Tg*(*dll4:Gal4FF*^*hu10049*^)^40^, Tg(UAS:tagRFP)^41^, *Tg*(*flt4: mCitrine*)^42^, *Tg*(*flt1enh:tdTomado*)^34^, *Tg*(*fli1a:lifeactCherry*)^43^, *Tg*(*EPV. Tp1-Mmu.Hbb:d2GFP*)^*mw4344*^ abbreviated as *Tg*(*Tp1:d2GFP*), *Tg*(*UAS:myc-Notch1a-intra*)^*kca3 45*^ *abbreviated as Tg*(*UAS:NICD*), *Tg*(*flk:DENDRA2*)^46^, *Tg*(*fli1a:B4GALT1-mCherry*)^bns9 17^.

### Morpholino and RNA injections

*Embryos were injected at the one-cell stage* with 1 nl morpholino oligonucleotides (MOs) or RNA. *Delta like 4 (dll4) splice-site* MO (GeneTools) (*5-TGATCTCTGATTGCTTACGTTCTTC-3)^34^ at* 1 ng/ nl; *gata1a* MO (5-CTGCAAGTGTAGTATTGAAGATGTC-3)^47^ 4ng/nl and a *tripartite motif containing 33 (trim33, a.k.a Tif1-γ) (5-* GCTCTCCGTACAATCTTGGCCTTTG-3)^48^ at 3 ng/ul. *Capped CAS9 mRNA was synthesized from linearized pCS2+ constructs using the mMessage mMachine SP6 kit (Ambion, AM1340), and was injected into embryos at a concentration of 200ng.* A gene specific oligo containing the SP6 promoter and *dll4* guide RNA sequence was annealed to the constant oligo as previously described^49^. *dll4* gRNA *5-tttgcctaaaaaactacc-3* was transcribed using the SP6 MEGAscript kit (Ambion, AM1330).

### Constructs

H2b-EGFP and H2b-mCherry in SIN18.hPGK.eGFP.WPRE lentiviral vector were gifts from J.H. Price(Kita-Matsuo et al., 2009)physiological and tissue engineering studies critical to the development of successful myocardial regeneration therapies require new ways to effectively visualize and isolate large numbers of fluorescently labeled, functional cardiomyocytes. Methodology/Principal Findings: Here we describe methods for the clonal expansion of engineered hESCs and make available a suite of lentiviral vectors for that combine Blasticidin, Neomycin and Puromycin resistance based drug selection of pure populations of stem cells and cardiomyocytes with ubiquitous or lineage-specific promoters that direct expression of fluorescent proteins to visualize and track cardiomyocytes and their progenitors. The phospho-glycerate kinase (PGK. GFP-α-tubulin in pRRL.PPT.CMV lentiviral vector was kindly provided by O. Pertz. Lentiviruses were produced as described previously^50^.

### Microscopy

Live microscopy was done in environmentally controlled microscopy systems based on a Nikon TE 2000 brightfield microscope^51^, a Perkin Elmer UltraView spinning disk confocal microscope, a Leica TSP LSM 5 confocal microscope, a Leica TCS SP8 DLS or a Zeiss LSM 880 Airyscan. For all imaging, embryos were embedded into an imaging chamber submerged in E3 medium containing 0.2% Ethyl 3-aminobenzoate methanesulfonate (MS222; sigma E10521) at a temperature of 28.5 °C^52^. Imaging was subsequently done with either 20x/0.75, 40x/1.4 or 63x/1.46 objectives. HUVECs and HUAECs cultured on fibronectin coated glass cover slips and application of microfluidic devices were done as previously described^18,51^. Tracking and polarization analysis of ECs *in vitro* was done using ImagePro 6.1 (MediaCybernatics, Bethesda, MD) and home-built applications in Matlab (Mathworks, Natick, MA).

### Microfluidic devices

Microfabrication of PDMS-based microfluidic devices and imaging chamber was done as previously described^51^.

### Cell Culture

*Culturing of ECs in vitro.* Culturing of HUVECs and HUAECs (Lonza, Basel, Switzerland) was done as previously described^51^.

### Statistical analysis

For each *in vivo* experiment, animals from the same clutch were divided into different treatment groups without any bias. The whole clutch was excluded if more than 10% of control embryos displayed obvious developmental defects. At least 20 animals from each treatment group were randomly picked for analysis. At least 3 independent experiments were performed per each treatment group. For *in vitro* experiments, at least 3 independent experiments were performed per each condition. Statistical analysis was performed using SPSS 20 (IBM). Mann-Whitney U test was used for statistical analysis of two groups, unequal variances. Unpaired t-test was used for two groups, equal variances. Kruskal-Wallis test was used for statistical analysis of multiple groups, equal variances, and 1-way ANOVA, for multiple groups, unequal variances. Dunns post-hoc test was used for pairwise multiple comparisons. P values < 0.05 were considered significant.

---

**Extended Data Figure 1:**
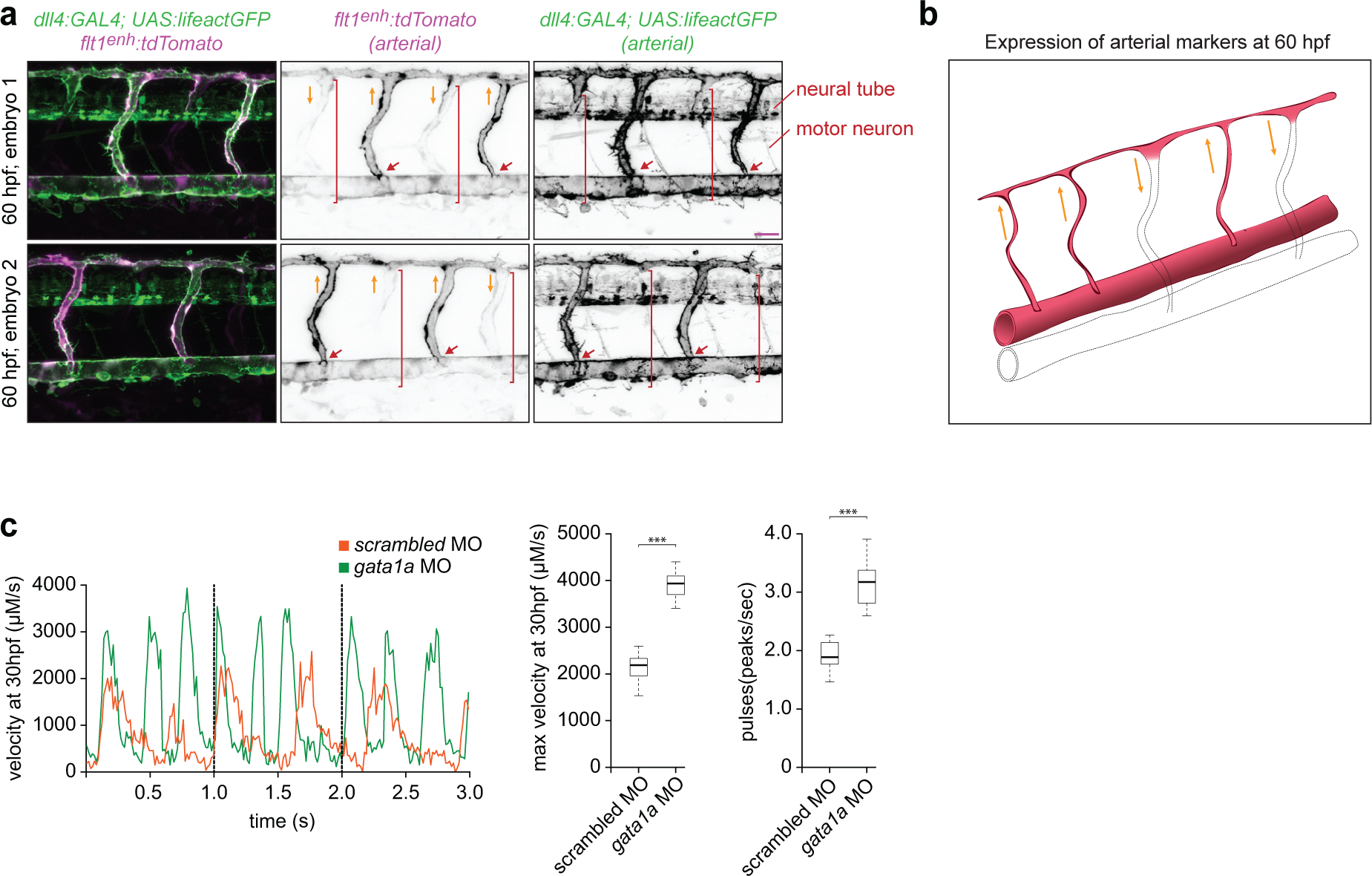
Imaging of arterial ECs in zebrafish embryo. **a**) *Tg(flt1*^*enh*^*:tdTomato*) and *Tg(dll4:GAL4; UAS:lifeactGFP*) transgenic lines that label arterial ECs. Lateral images of zebrafish embryos at 60 hpf with anterior side facing left. *Orange arrows* indicate the direction of blood flow, *red arrows* point to arterial ISVs and *brackets* highlight regions of venous ISVs without arterial ECs. Note that, in addition to arteries, tdTomato is also weakly expressed in veins, whereas lifeactGFP is not detectable in veins but expressed in the neural tube and motor neurons. Scale bar is 25 μm. **b**) Schematic of the spatial expression of arterial markers. **c**) Blood velocity in the DA embryos with and without erythrocytes. *Left panel* shows velocity as a function of time. *Middle panel* shows maximal velocity. *Right panel* shows box plot of the pulses.

**Extended Data Figure 2:**
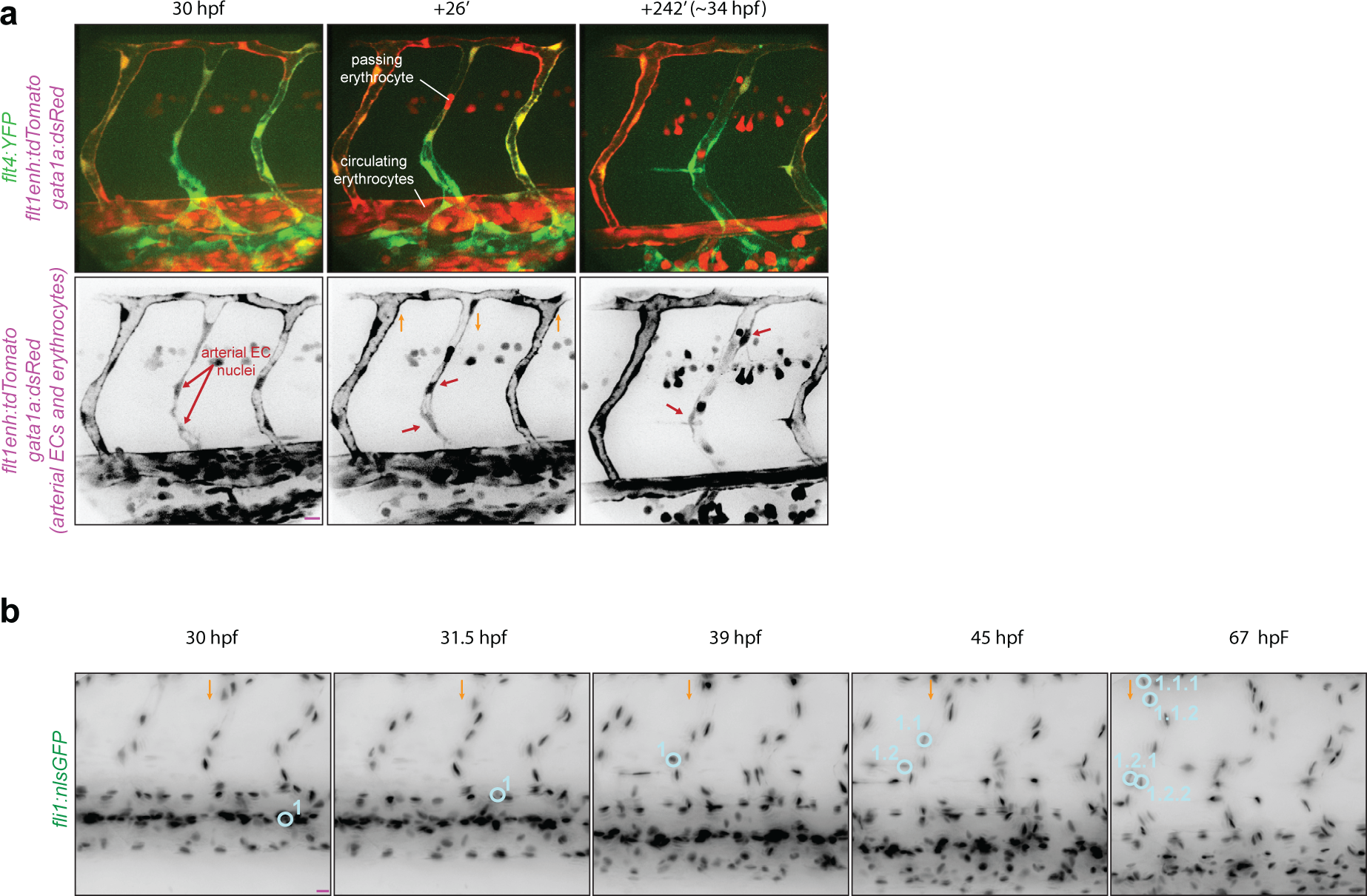
Blood flow-dependent displacement of arterial ECs by venous ECs in venous ISVs. **a**) Lateral images of zebrafish embryos with anterior side facing left. **b**) Stills from Video 1. Venous ECs are labelled with mCitrine, arterial ECs are labelled with mCitrine and tdTomato, and erythrocytes are labelled with dsRed. **c**) Stills from Video 3. Proliferation of venous ECs in a venous ISV. Orange arrows indicate the direction of blood flow through the ISVs. Scale bars are 25 μm.

**Extended Data Figure 3:**
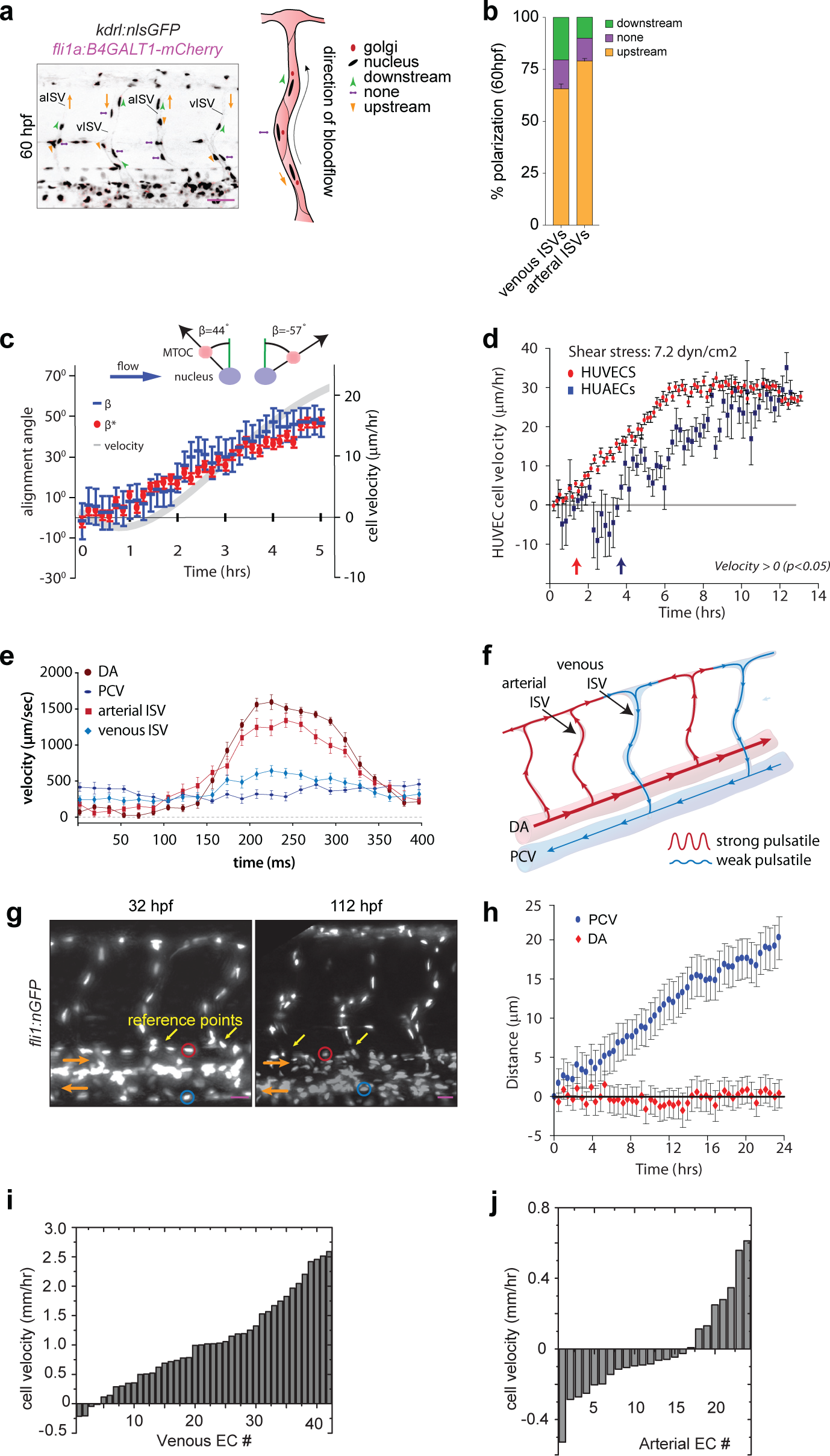
Blood flow promotes EC migration in veins but not in arteries. Panels in (**a**) and (**g**) show lateral images of zebrafish embryos with anterior side facing left. *Orange arrows* indicate the direction of blood flow. Error bars are SEM. **a**) All ECs express nuclear-GFP and mCherry-fused marker of the Golgi. Planar polarization of ECs in ISVs is measured by the vector connecting the nucleus with the Golgi. Scale bar is 25 μm. **b**) Quantification of EC planar polarization in venous and arterial ISVs (n=10 embryos). **c**) Positions of microtubule organization complexes (MTOCs) and nuclei in individual HUVECs and their instantaneous velocities were monitored for 300 min after the exposure to flow with a shear stress of 7.2 dyn/ cm^2^. *Blue dashes* show the values of the polarization angle, β, with 90° corresponding to polarization against the flow and β = -90° - polarization along the flow. *Red circles* show the values of the migration angle, β^*^, with 90° corresponding to migration against the flow and β^*^ = -90° -migration along the flow. *Grey line* (ordinate on the right) show the average cell migration velocity in the upstream direction. **d**) Average velocities of upstream migration as functions of time after the inception of shear flow (n=250). *Arrows* at the bottom (colors correspond to those of the velocity data points) indicate the time points at which the migration of cells against the flow becomes statistically significant (average upstream velocity becomes positive with p<0.05). **e**) Velocity of erythrocytes as a function of time within one heart beat (n=16 vessels per condition; average of large number of heart beats per vessel). **f**) Schematic representation of the dynamics of blood flow in the intersegmental vasculature. **g**) All ECs express nuclear-localized GFP. Intersegmental vessels (*yellow arrows*) serve as reference points for determining the location of tracked arterial (*red ovals*) and venous (*blue ovals*) ECs. Scale bar is 30 μm. **h**) Average displacements of ECs in the PCV (*blue*, n=42, 6 embryos) and DA (*red*, n=25, 6 embryos) as functions of time. Displacement of ECs was analyzed with at least 6 venous and 4 arterial ECs per embryo. Displacement is considered positive, if the cell migrates upstream. **i,j**) Histograms of velocities of venous (i) and arterial (j) ECs. Velocities are averaged over the course of experiment and ordered by their values. Velocity is considered positive, if the cell migrates upstream.

**Extended Data Figure 4:**
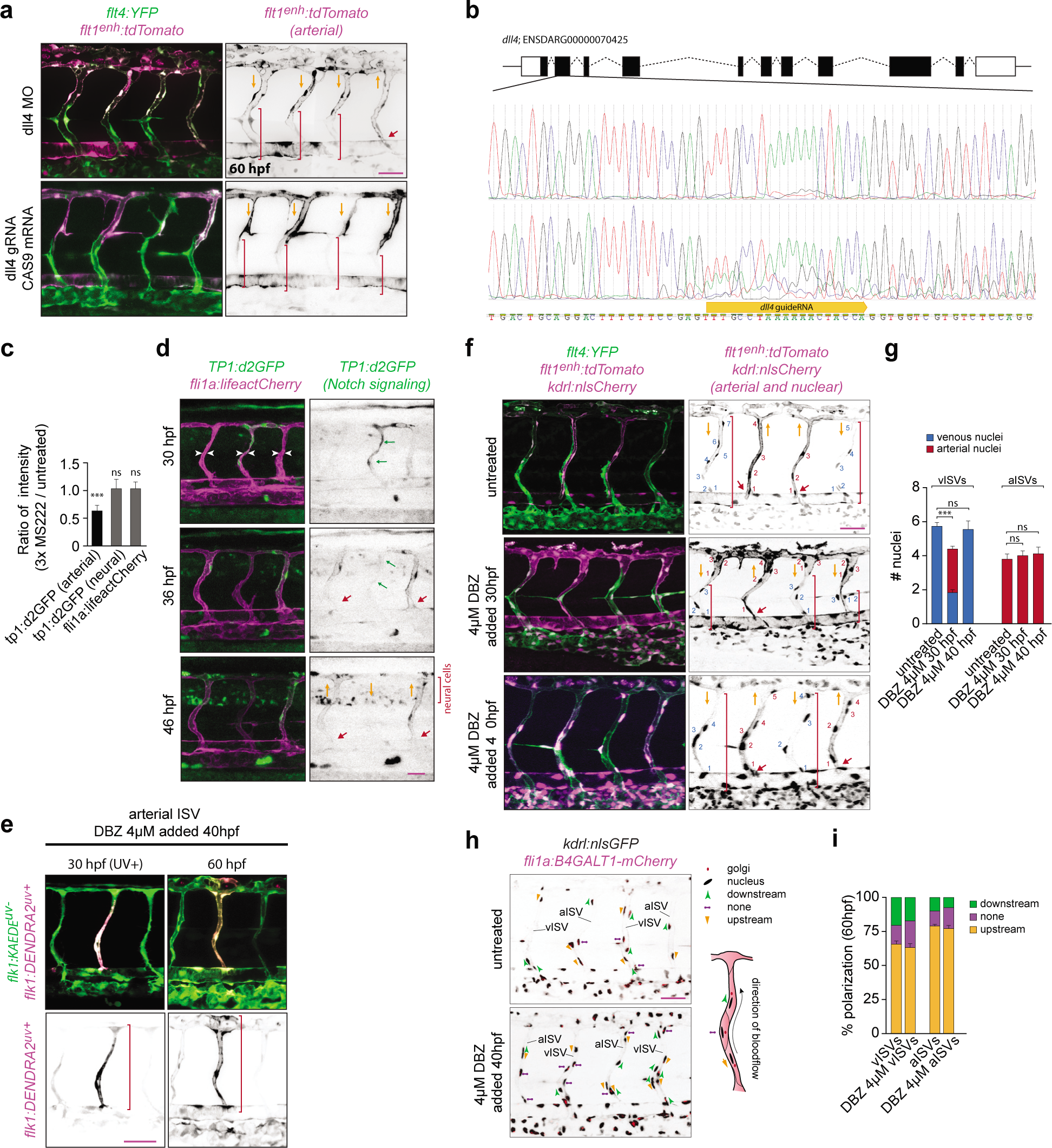
Arterial blood flow protects ISVs from transforming into veins by activating Notch signaling. Panels in (**a**), (**d**), (**g**), (**f**) and (**i**) show lateral images of zebrafish embryos with anterior side facing left. Images for panels (**a**), (**g**) and (**i**) were acquired at 60 hpf. *Red arrows* point to arterial ISVs and *red brackets* highlight regions of venous ISVs without arterial ECs. *Orange arrows* indicate the direction of blood flow. Scale bars are 25 μm. All numbers are averages ± SEM from at least three independent experiments with a minimum of n=25 animals per conditions per experiment. ‘***’ indicates P<0.001. **a**) Abrogation of *dll4* by MO or CRISPR-CAS9. Venous ECs are labelled with mCitrine and arterial ECs are labelled with mCitrine and tdTomato. **b**) CRISPR-CAS9 targeted location and representative sequencing plot of an injected embryo. **c**) ECs express lifeact-mCherry. Destabilized GFP (d2GFP) is expressed under the control of 12xCSL Notch responsive elements. *White arrowheads* highlight ISVs without blood flow. *Green arrows* highlight an ISV with above-the-average expression of d2GFP prior to initiation of blood flow. **d**) Quantification of the effect of a reduction in blood flow on d2GFP expression in arterial ISVs (n=10 embryos per condition; 4-6 ISVs per embryo). **e**) The inhibition of Notch does not have affect EC migration in an arterial ISV. All ECs express the photoconvertible (green-to-red) fluorescent protein *DENDRA2. Red brackets* highlight ECs with photo-converted DENDRA2. The photo-converted was done at 30 hpf in an arterial ISV. **f**) The effect of inhibition of Notch on the number of ECs in ISVs. Arterial ECs express mCitrine, tdTomato and nuclear-Cherry, and venous ECs express mCitrine and nuclear-Cherry. *Red brackets* depict venous ECs in venous ISVs. **g**) Number of arterial and venous ECs in arterial and venous ISVs in experiments with inhibition of Notch. **h**) Notch inhibition does not affect EC polarization in ISVs. All ECs express nuclear-GFP and mCherry-fused marker of the Golgi. Planar polarization of ECs in ISVs is measured by a vector connecting the nucleus with Golgi. **i**) Quantification of EC planar polarization in venous and arterial ISVs in experiments with inhibition of Notch.

## Supplementary Videos

**Supplementary Video 1. Upstream migration of arterial ECs in a venous ISV.** Time-lapse imaging of *Tg(flt4:mCitrine; flt1*^*enh*^*:tdTomato; gata1a:dsRed*) in which venous ECs are labelled with mCitrine (*green*), arterial ECs are labelled with mCitrine and tdTomato, and erythrocytes are labelled with dsRed (red). *Orange arrows* depict first erythrocyte passing through venous ISVs. *Green arrow* depicts arterial ECs migrating upstream in a venous ISV. *Red arrow* points to arterial ISVs.

**Supplementary Video 2. Displacement of arterial ECs by venous ECs in a venous ISV.** Timelapse imaging of *Tg(dll4:GAL4; UAS.lifeactGFP; flila.lifeactCherry*) in which venous ECs express lifeactCherry (*magenta*) and arterial ECs express lifeactCherry and lifeactGFP (*green). Red arrow* points to an arterial EC migrating dorsally in a venous ISV.

**Supplementary Video 3. Migration and proliferation of venous ECs in a venous ISV.** Time-lapse imaging and cell tracing in *Tg(kdl:H2B-Cherry*), in which all ECs express nuclear-localized mCherry (*grey*).

**Supplementary Video 4. Arterial ECs do not migrate in an arterial ISV.** Time-lapse imaging of *Tg(dll4:GAL4; UAS.lifeactGFP; flila.lifeactCherry*) in which venous ECs express lifeactCherry (*magenta*) and arterial ECs express lifeactCherry and lifeactGFP (*green). Red arrow* points to an arterial EC in an arterial ISV.

**Supplementary Video 5. Flow-induced changes in the planar polarization and migration of HUVECs in the microfluidic model.** Positions of MTOC and nuclei in individual HUVECs were monitored for 300 min after their exposure to flow with a shear stress τ = 7.2 dyn/cm^2^. The direction of flow is from left to right. *Left panel shows fluorescence images* (*with an inverted grayscale*) of a fragment of a microchannel with ECs expressing GFP-α-tubulin. The arrows show the direction of cell polarization and their color indicates the polarization angle, β, with red corresponding to 90° (polarization against the flow) and blue corresponding to β = -90° (polarization along the flow). *Right panel* shows phase contrast images of the same area. The arrows show the direction of cell migration and their color represents the angle between the directions of migration and flow, β^*^, with red corresponding to 90° (migration against the flow) and blue corresponding to β^*^ = -90° (migration along the flow). The dials in the top left corners show color-coded polarization angle, β (*left panel*), and migration angle, β^*^ (*rightpanel*) averaged over all cells in the field of view. Images were acquired every 10 min. Frame rate is 3 per sec (1 sec = 30 min). Scale bar is 30 μm.

**Supplementary Video 6. Migration of HUVECs exposed to low and high shear stress in the microfluidic model.** Phase-contrast images of HUVECs exposed to a shear stress of 0.23 dyn/cm^2^ (*upper panel*) and 14.5 (*lower panel*) acquired every 10 min. Direction of the flow was from left to right. Frame rate is 6 per sec. Scale bar is 50 μm.

**Supplementary Video 7. Blood flow in arterial and venous ISVs.** Time-lapse imaging of *Tg(gata1:dsRed*), in which all erythrocytes express dsRed (*grey*).

**Supplementary Video 8. Blood flow in the dorsal aorta and posterior cardinal vein.** Time-lapse imaging of *Tg*(*gata1:dsRed*), in which all erythrocytes express dsRed (*grey).*

**Supplementary Video 9. Expression of *Tp1:d2GFP* during remodeling of the ISVs.** Time-lapse imaging of *Tg*(*Tp1:d2GFP; flila.lifeactCherry*) which expresses a de-stabilized GFP (*green*) under the control of 12xCSL Notch responsive elements. All ECs are marked by lifeactCherry (*magenta*). *Red arrows* point to arterial ISVs. *Green arrow* depicts a venous sprout that did not anastomose with an arterial ISV with high Notch activity.

**Supplementary Video 10. Expression of *Tp1:d2GFP* during remodeling of the ISVs.** Time-lapse imaging of *Tg(Tp1:d2GFP; fli1a:lifeactCherry*) which expresses a de-stabilized GFP (*green*) under the control of 12xCSL Notch responsive elements. All ECs are marked by lifeactCherry (*magenta). Red arrows* point to arterial ISVs. *Green arrows* depict venous sprouts. The sprout on the *left* anastomoses with an ISV. *Orange arrow* highlights the formation of a lumen in the sprout on the right. This sprout detaches from an ISV aft er d2GFP becomes expressed. *Blue arrow* points to a venous EC migrating from the sprout into this ISV.

**Supplementary Video 11. Expression of *Tp1:d2GFP* during remodeling of the ISVs.** Time-lapse imaging of *Tg(Tp1:d2GFP; fli1a:lifeactCherry*) which expresses a de-stabilized GFP (*green*) under the control of 12xCSL Notch responsive elements. All ECs express lifeactCherry (*magenta). Red arrows* point to arterial ISVs. *Green arrows* point to ECs expressing d2GFP in venous ISV.

## ACKNOWLEGMENTS

We thank M. Ginsberg, S. Schulte-Merker and A. van Impel for comments on the manuscript; D. Stainier for the golgi reporter line; S. Schulte-Merker for the Dll4, Flt1enh and Flt4 reporter lines. N. Chi for the dendra line. P. Tsai for use of the Q-bio confocal. This work was supported by NIH grants HL124195 to ET, BW, AG, EG, DT and HL078784 to ET; AHA award 14SDG20380181 to ET; EMBO fellowship ALTF_757-2014 to BW; and NSF1411313 grant to AG and EG.

## AUTHOR CONTRIBUTIONS

ET and BW designed and performed the experiments with zebrafish. EG, AG, and ET designed experiments in microfluidic devices. BW and ET performed microfluidic experiments. SKS, BW, and ET analyzed the data. BW, AG, and ET wrote the manuscript. DT edited the manuscript and reviewed the data.

